# Multiple evolutionary origins and losses of tooth complexity in squamates

**DOI:** 10.1101/2020.04.15.042796

**Authors:** Fabien Lafuma, Ian J. Corfe, Julien Clavel, Nicolas Di-Poï

**Affiliations:** Developmental Biology Program, Institute of Biotechnology, University of Helsinki, FIN-00014 Helsinki, Finland; Department of Life Sciences, The Natural History Museum, London SW7 5DB, United Kingdom; Laboratoire d’Écologie des Hydrosystèmes Naturels et Anthropisés (LEHNA), Université Claude Bernard Lyon 1 – UMR CNRS 5023, ENTPE, F-69622 Villeurbanne, France

## Abstract

Teeth act as tools for acquiring and processing food and so hold a prominent role in vertebrate evolution^1,2^. In mammals, dental-dietary adaptations rely on tooth shape and complexity variations controlled by cusp number and pattern – the main features of the tooth surface^3,4^. Complexity increase through cusp addition has dominated the diversification of many mammal groups^3,5-9^. However, studies of Mammalia alone don’t allow identification of patterns of tooth complexity conserved throughout vertebrate evolution. Here, we use morphometric and phylogenetic comparative methods across fossil and extant squamates (“lizards” and snakes) to show they also repeatedly evolved increasingly complex teeth, but with more flexibility than mammals. Since the Late Jurassic, six major squamate groups independently evolved multiple-cusped teeth from a single-cusped common ancestor. Unlike mammals^10,11^, reversals to lower cusp numbers were frequent in squamates, with varied multiple-cusped morphologies in several groups resulting in heterogenous evolutionary rates. Squamate tooth complexity evolved in correlation with dietary change – increased plant consumption typically followed tooth complexity increases, and the major increases in speciation rate in squamate evolutionary history are associated with such changes. The evolution of complex teeth played a critical role in vertebrate evolution outside Mammalia, with squamates exemplifying a more labile system of dental- dietary evolution.

---

As organs directly interacting with the environment, teeth are central to the acquisition and processing of food, determine the achievable dietary range of vertebrates^1^, and their shapes are subject to intense natural selective pressures^8,12^. Simple conical to bladed teeth generally identify faunivorous vertebrates, while higher dental complexity – typically a result of more numerous cusps – enables the reduction of fibrous plant tissue and is crucial to the feeding apparatus in many herbivores^4,8,13^. Evidence of such dental-dietary adaptations dates back to the first herbivorous tetrapods in the Palaeozoic, about 300 million years ago (Ma)^13^. Plant consumers with highly complex teeth have subsequently emerged repeatedly within early synapsids^13^, crocodyliforms^14^, dinosaurs^15^, and stem- and crown-mammals^6,7,9,16^. Since the earliest tetrapods had simple, single-cusped teeth^8^, such examples highlight repeated, independent increases of phenotypic complexity throughout evolution^17^. Many such linked increases in tooth complexity and plant consumption have been hypothesised to be key to adaptive radiations^6,9^, though such links have rarely been formally tested. It is also unclear whether the known differences in tooth development between tetrapod clades might result in differences in the evolutionary patterns of convergent functional adaptations^18,19^.

To understand the repeated origin of dental-dietary adaptations and their role in vertebrate evolution, we investigated tooth complexity evolution in squamates, the largest tetrapod radiation. Squamata is recognized for including species bearing complex multicuspid teeth within heterodont dentitions^20^, and squamate ecology spans a broad range of past and present niches. Squamates express dental marker genes broadly conserved across vertebrates^18^, with varying patterns of localization and expression compared to mammals, and structures at least partially homologous to mammalian enamel knots (non-proliferative signalling centres of ectodermal cells) determine tooth shape in some squamate clades^19,21,22^. In mammals – the most commonly studied dental system – novel morphologies arise from developmental changes in tooth morphogenesis^23^. Epithelial signalling centres – the enamel knots – control tooth crown morphogenesis^24^, including cusp number and position and ultimately tooth complexity, by expressing genes of widely conserved signalling pathways^18,25^. Experimental data show most changes in these pathways result in tooth complexity reduction, or complete loss of teeth^25^, yet increasing tooth complexity largely dominates the evolutionary history of mammals^6-9,16^. To determine whether similar patterns of tooth complexity underlie all tetrapod evolution or are the specific results of mammalian dental development and history, we used morphometric and phylogenetic comparative methods with squamate tooth and diet data.

We analysed cusp number and diet data for 545 squamate species spanning all living and extinct diversity and identified species with multicuspid teeth in 29 of 100 recognized squamate families (Figure 1a & Extended Data Figure 1). Within extant “lizards”, we found multicuspid species in almost 56% of families (24/43). While lacking entirely in mostly predatory clades (dibamids, geckos, snakes), multicuspidness dominates Iguania and Lacertoidea, the two most prominent groups of plant-eating squamates. A Kruskal–Wallis test and post-hoc pairwise Wilcoxon–Mann–Whitney tests show squamate dietary guilds differ statistically in tooth complexity, with the proportion of multicuspid species and cusp number successively increasing along a plant consumption gradient, from carnivores to insectivores, omnivores, and herbivores (*p*-value < 0.001; Fig. 1b, Extended Data Table 1). We quantified tooth outline shape in a subset of taxa spanning all major multicuspid groups with two-dimensional semi-landmarks, which showed the teeth of herbivores are more protruding with a wider top cusp angle (Fig. 1c). A regularized phylogenetic multivariate analysis of variance (MANOVA) on principal component scores confirm statistically significant differences between diets overall (*p*-value = 0.001; Fig.1c) with negligible phylogenetic signal in the model’s residuals (Pagel’s λ = 0.03). Herbivore teeth differ from both the insectivorous and omnivorous morphospace regions (Fig.1c, Extended Data Table 2), similarly to observations from mammals and saurians^4,20^. Furthermore, we find support for shifts in the rate of evolution of tooth shape outline independent of cusp number among the 75 species examined (log Bayes Factor = 319), with particularly high rates characterising Iguanidae (Extended Data Figure 2).

**Fig. 1.**
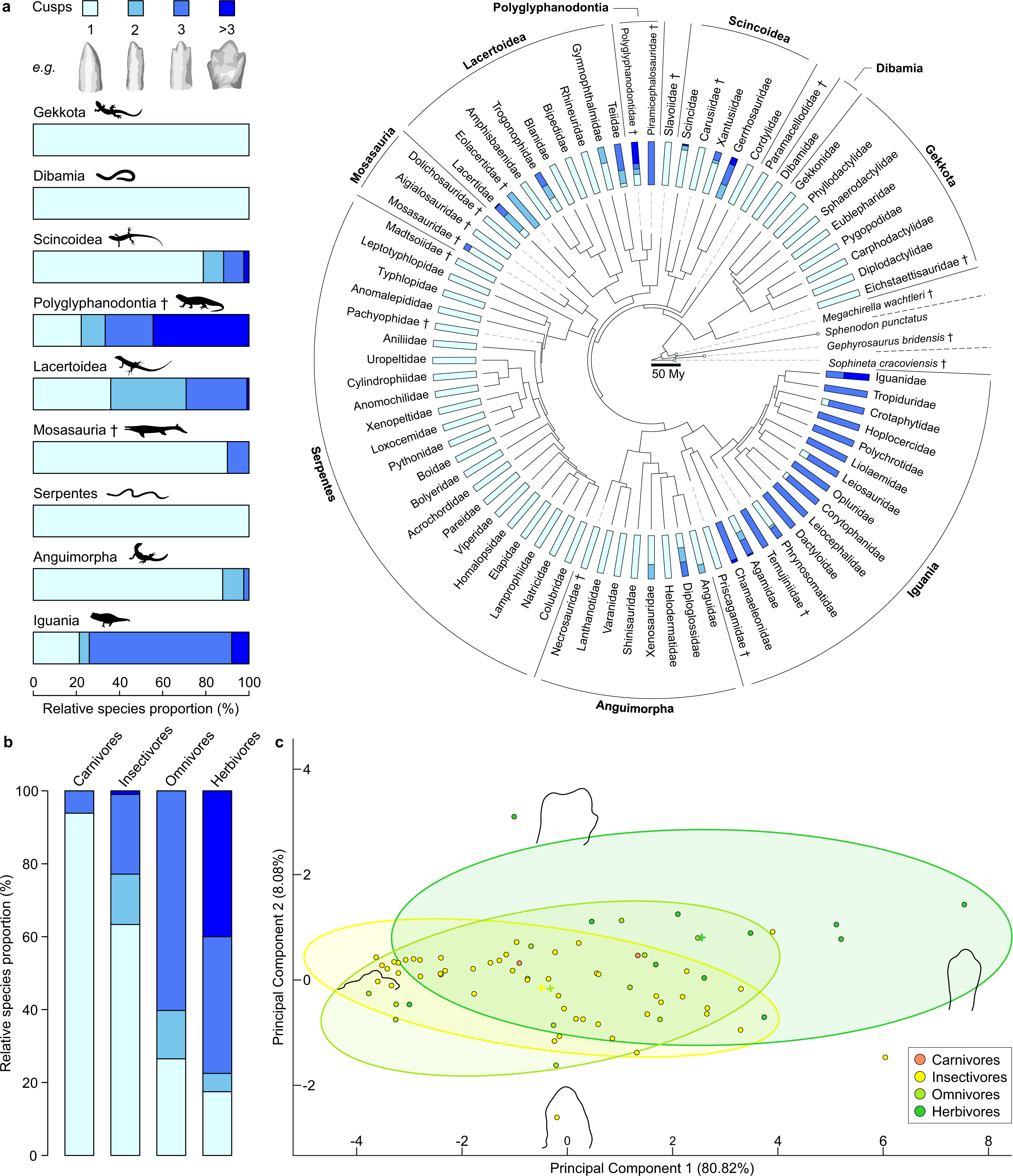
The diversity of squamate dental morphologies correlates with a gradient of plant consumption. **a**, Relative proportions (%) of tooth complexity levels in all known squamate suborders/super-families (left) and 76 families (right) based on cusp number data for 545 living and extinct species (including the most ancient known squamate *Megachirella wachtleri*), two rhynchocephalians, and the stem-lepidosaurian *Sophineta cracoviensis*, with example teeth for each complexity level redrawn from microCT-scan data (not to scale). **b**, Relative proportions (%) of tooth complexity levels in 545 squamates sorted by diet. **c**, Discrete Cosine Transform analysis of multicuspid tooth labial view profiles from 75 extant and fossil squamate species, with 95% confidence ellipses for insectivorous, omnivorous, and herbivorous species. Theoretical tooth profiles at the extreme positive and negative values of each axis reconstructed from the first 21 harmonic coefficients. Scalebar = 50 million years (My). † = extinct taxon. Silhouettes: the authors, Phylopic, and public domain (see Methods for license information).

Using Maximum-Likelihood reconstructions of ancestral character states^26^ across our squamate phylogeny (Fig. 2, Extended Data Figure 3 and 4, Supplementary Table 1 and 2), we found dental-dietary adaptations to plant consumption repeatedly evolved, arising from the convergent evolution of multicuspidness. Since the Late Jurassic, six major clades and 18 isolated lineages independently evolved multicuspid teeth from a unicuspid ancestral morphology, mostly through single-cusp addition events. Similar numbers – 10 clades, 13 isolated lineages – show independent origins of plant consumption from carnivorous or insectivorous ancestors (Fig. 2, Supplementary Table 3). Across the tree, most lineages and terminal taxa are unicuspid insectivores retaining the reconstructed ancestral squamate condition. However, of 102 lineages showing cusp number or plant consumption increases, 42 (41%) of increases are along the same phylogenetic path than an increase in the other character (see Methods).

**Fig. 2.**
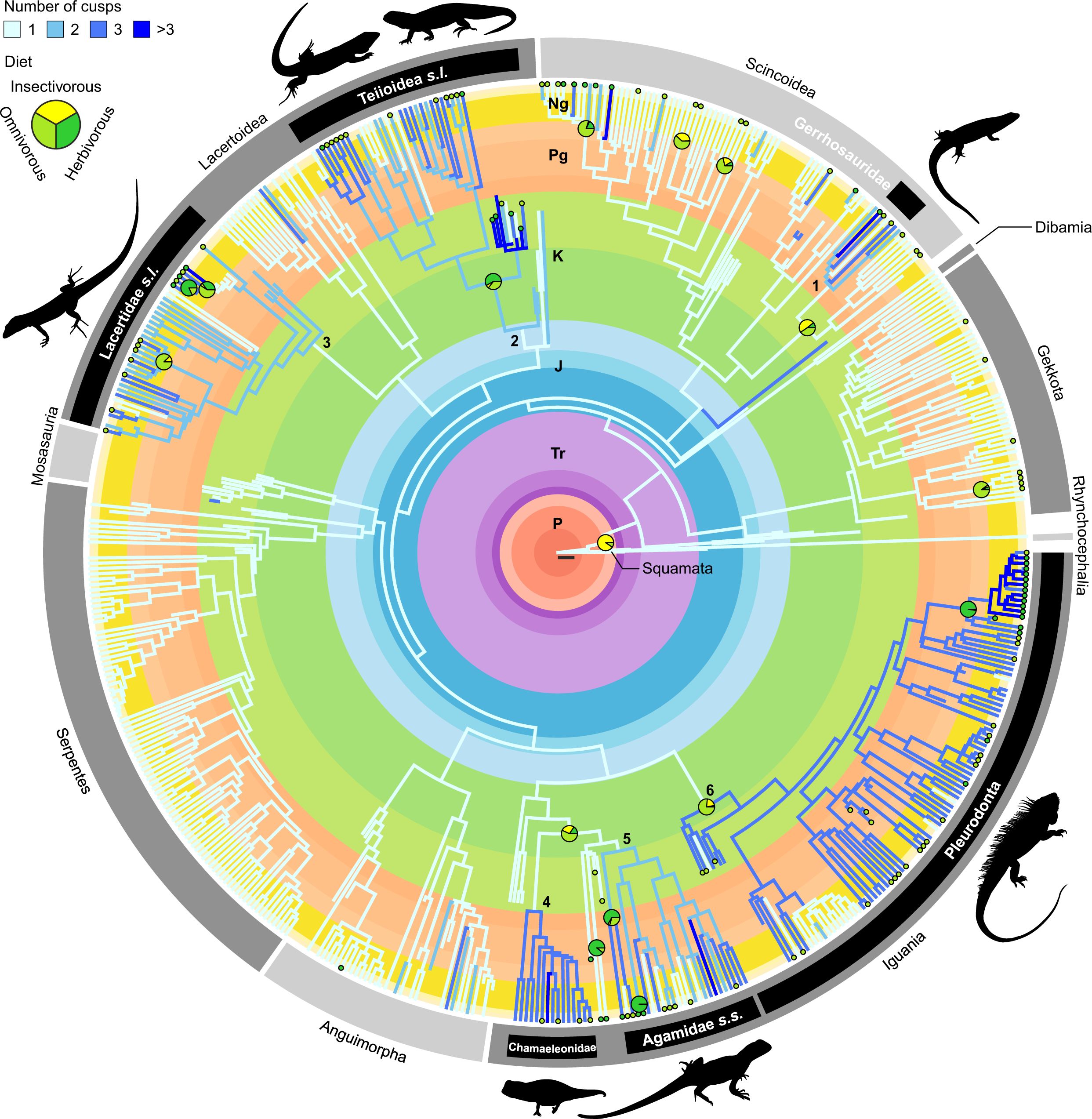
Multiple independent acquisitions of multicuspid teeth and plant consumption are found across the squamate phylogeny. Known and Maximum-Likelihood ancestral state reconstructions of the number of cusps (branch colours) and diet (node pie charts and branch tip small circles) in squamates. Pie charts indicate the most ancient nodes with >50% combined relative likelihood for omnivorous and herbivorous diets; also shown are the first nodes with >50% relative likelihood for herbivory within already omnivorous clades. Branch tip circles indicate omnivorous/herbivorous species. Six major clades showing independent originations of multicuspid teeth – 1: Gerrhosauridae. 2: Teiioidea + Polyglyphanodontia (informally Teiioidea *sensu lato*). 3: total group Lacertidae (informally Lacertidae *sensu lato*). 4: Chamaeleonidae. 5: non-Uromastycinae agamids (informally Agamidae *sensu stricto*). 6: total group Pleurodonta. P: Permian. Tr: Triassic. J: Jurassic. K: Cretaceous. Pg: Paleogene. Ng: Neogene. Scalebar = 10 million years. Silhouettes: the authors, Phylopic, and public domain (see Methods for license information).

Squamate dental evolution was however labile and included repeated reversals towards lower tooth complexity. Both complexity and diet changed similarly through much of squamate evolution, though there were more changes in diet than complexity (115 vs. 92 lineages), and reversals in diet were more common than for complexity (56% vs. 44%) (Fig. 3, Extended Data Figure 3). Such flexibility is reflected in the reconstructed transition rates underlying our models of evolution for tooth complexity and diet, where higher relative rates characterise decreases in cusp number and plant consumption compared to increases (Extended Data Figure 4, Supplementary Table 1 and 2). Moreover, 38% of inferred complexity decreases were due to the simultaneous loss of two cusps or more, while multiple-cusp addition events were half (20%) as frequent. We identify two lineages (genera *Gallotia* and *Phrynosoma*) in which multicuspid teeth re-evolved following earlier loss (Fig. 2, Extended Data Figure 3). Most often, reversals to lesser cusp numbers followed a decrease in plant consumption (52% of paired events; Supplementary Table 3), likely resulting from the relaxation of selective pressures for plant consumption.

**Fig. 3.**
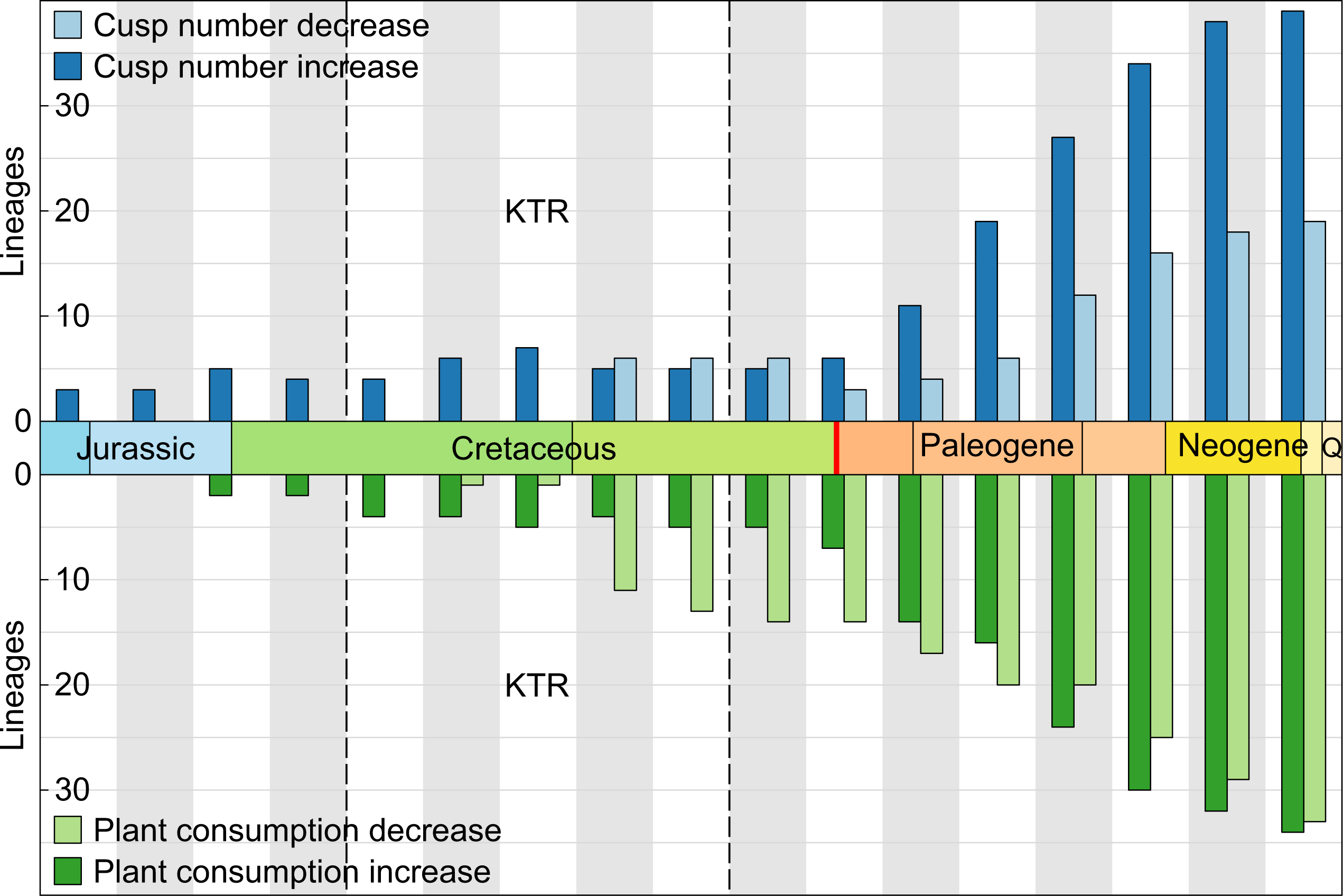
Dynamics of squamate tooth complexity and plant consumption evolution over the last 165 million years. Lineages showing increasing (n = 61) or decreasing (n = 31) tooth complexity (top) and increasing (n = 51) or decreasing (n = 64) plant consumption (bottom) per ten million year-time bins. KTR: Cretaceous Terrestrial Revolution (125–80 Ma). Q: Quaternary. Decreases in both cusp number and plant consumption proportion first outnumber increases during the Cretaceous Terrestrial Revolution (KTR), while the Cretaceous–Paleogene boundary (in red) shows the change towards the Cenozoic pattern of approximately twice as many cusp increases as decreases, and similar numbers of plant consumption increase and decrease from the Paleogene on.

Furthermore, we find the observed dental-dietary patterns derive from the correlated evolution of tooth complexity and plant-based diets under highly variable rates of phenotypic evolution. Our results show strong support for a correlated model of the evolution of multicuspidness and plant consumption, which assumes transition rates in one trait directly depend on character state in the other trait (log Bayes Factor = 21, Supplementary Table 4). Additionally, a model with heterogenous character transition rates throughout the tree better fits the macroevolutionary pattern of each trait than a constant model, with the highest rates observed resulting in a relatively balanced mixture of tooth complexity and diet character states for the clades concerned (*e.g.*, Lacertidae) (Figs. 2, 4a, 4b, Supplementary Table 5).

**Fig. 4.**
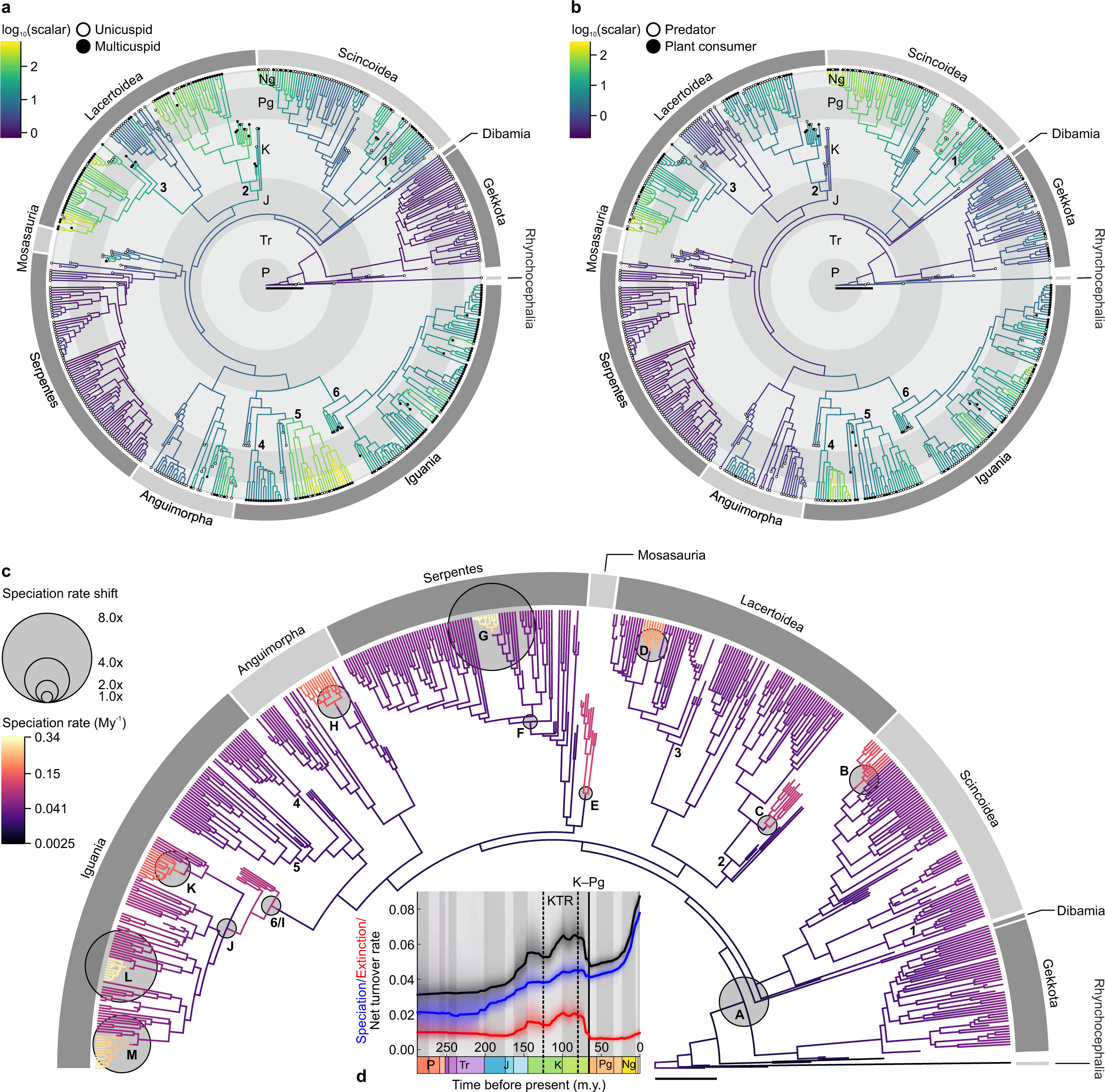
Dental-dietary rates of phenotypic evolution and squamate macroevolution. Log-transformed averaged rate scalars of the character transition rates of tooth complexity (**a**) and diet (**b**) across squamates. Positive values (*i.e.*, rate scalar > 1) indicate increased relative transition rates. **c**, Rates of squamate speciation for one maximum shift credibility configuration (MSC) out of ten similar independent replicates. 13 rate shifts (A–M) present in at least five MSC replicates are indicated proportionally to their magnitude compared to the background rate. **d**, Mean rates of squamate speciation, extinction, and net turnover through time (in My^-1^). 1: Gerrhosauridae. 2: Teiioidea + Polyglyphanodontia (informally Teiioidea *sensu lato*). 3: total group Lacertidae (informally Lacertidae *sensu lato*). 4: Chamaeleonidae. 5: non-Uromastycinae agamids (informally Agamidae *sensu stricto*). 6: total group Pleurodonta. P: Permian. Tr: Triassic. J: Jurassic. K: Cretaceous. Pg: Paleogene. Ng: Neogene. Q: Quaternary. KTR: Cretaceous Terrestrial Revolution (125–80 Ma). K–Pg: Cretaceous–Paleogene extinction event (66 Ma). Scalebars = 50 million years.

These evolutionary increases in tooth complexity and plant consumption appear to have contributed to the diversification of Squamata. Using models with variable rates of diversification implemented in a Bayesian framework (through a reversible jump Markov Chain Monte Carlo algorithm) and allowing for the inclusion of fossil taxa, we identified multiple events of increased speciation (13 events in the studied tree with up to eight-fold magnitude for the focal group versus its outgroup; Fig. 4c, Extended Data Table 3). Five speciation increases coincide exactly with increases in tooth complexity or plant consumption, and three more are just one node away from such increases. The equivalent results for decreases are two and one lineage respectively (Extended Data Table 3, Supplementary Table 6). We further tested this apparent association between diversification shifts and transitions of cusp number and diets using a “hidden state” trait-dependent model of speciation and extinction. Results from this model suggest each trait (tooth complexity and diet) contributed considerably to the group’s diversification – with rates of speciation and extinction increasing with transitions to multicuspidness or plant-based diets – despite the influence of unobserved factors beyond our study (Extended Data Table 4 and Figure 5). Combining these results, we propose plant consumption and tooth complexity changes – principally increases – were critical innovations for squamate evolution.

The evolution of tooth complexity in Squamata encompasses multiple independent radiations defined by increasingly complex teeth. This mirrors patterns of mammalian diversification, in which stem-mammals show repeated independent evolutions of multicuspid teeth through the Palaeozoic and Mesozoic^7,9,13^, the key adaptations of tribosphenic or pseudo-tribosphenic molars separately originated in the Jurassic^27^, and quadritubercular molars with a hypocone appeared multiple times in the Cenozoic^3,5,6^. It differs from the mammalian pattern, however, in that the most recent common ancestor (MRCA) of Mammalia was multicuspid^7^, whereas we reconstruct the MRCA of Squamata as unicuspid and infer at least 24 independent acquisitions of multicuspidness in squamate lineages. Squamate tooth evolution was also not mainly unidirectional as in mammals, with numerous lineages losing tooth complexity, including reversals to the ancestral unicuspid condition. Moreover, tooth complexity at times subsequently re-emerged within lineages that previously underwent such reversals, in opposition to Dollo’s law^28^. Despite the lack of a similar large-scale phylogenetic assessment, studies suggest relatively few mammalian lineages experienced reversals towards reduced tooth complexity (including complete tooth loss)^10,11,29^, and even fewer re-evolved cusps once lost^30^. We confirm here across the whole of Squamata the link noted previously between plant-eating squamates and a specialized, typically more complex dentition^20^, similar to those hypothesized or discovered for early tetrapods^13^, crocodyliforms^14^, and mammals^4^. The generality of these findings suggests similar ecological and dietary selective pressures for complex dental phenotypes operate across all tetrapods. We find strong support for correlated evolution of multicuspidness and plant consumption, both of which promoted increased diversification of several major squamate groups (*e.g.*, Pleurodonta, Polyglyphanodontia), and propose environmental factors such as the floral turnovers of the Cretaceous Terrestrial Revolution (KTR – 125–80 Ma)^31,32^ and Cenozoic^33^ are the most plausible drivers of increasing plant consumption in squamate evolution^34^. During the KTR, squamate speciation locally peaked, net turnover was highest until the Late Miocene, and extinction was overall highest. All three metrics drop prior to and across the Cretaceous-Paleogene (K-Pg) boundary, suggesting the K-Pg extinction event had less of an effect on squamate diversification than the KTR. These KTR diversification shifts coincide with the majority of the only period where reductions in both tooth complexity and plant consumption outnumber increases, suggesting a rapidly shifting set of available dietary niches as previously proposed for mammals of the same time^35^. Reductions of squamate cusp number most often followed plant consumption reductions, suggesting relaxed selective pressures on diet enabled the loss of tooth complexity, as with mammals^11,29^. However, selective pressures on squamate teeth may not be as intense as for mammals. Most plant-eating squamates still consume insects^36^, suggesting that, unlike in Mammalia, no hyper-specialist ratchet operated^37,38^.

The patterns of squamate dental complexity evolution we observe offer a valuable counterpoint to the mammalian picture, exemplifying dental-dietary adaptations that responded to similar selective pressures, while resulting in more labile dental complexity throughout evolution. Despite vertebrates sharing a basic tooth gene-network^18^, mammal teeth are more integrated structures, less prone, through intense selective pressures, to loss of complexity, though also capable of accumulating significantly more variance and reaching farther phenotypic extremes over time^39^. Since such finely tuned dental morphologies and precise occlusion have a critical role in ensuring mammals meet their high energy needs^8^, endothermy may limit the possibilities of mammalian dental simplification compared to ectothermic squamates. Several dental developmental differences to mammals can be suggested to explain why squamates didn’t fall into a developmental complexity trap^40^, but instead evolved complex teeth highly liable to developmental instability and simplification^41^. These include simpler, less compartmentalised expression of dental development genes during tooth formation^19,22^, a less complex morphological starting point than mammal teeth^7^, and potentially simpler and/or looser gene regulatory networks^18^. We propose these characteristics of squamates explain both the evolutionary lability of their dental complexity and diet, and the near-complete absence of mammal-like teeth in over 250 million years of squamate history^42^.

## Methods

### Phylogenies

We gathered our own observations and reports from the literature on cusp number and diet for 548 species (429 extant species and 119 fossil species equally distributed between Mesozoic and Cenozoic). The data include all major squamate groups plus squamate stem taxa – including the oldest known squamate *Megachirella wachtleri*, two rhynchocephalians (the extant *Sphenodon punctatus* and the fossil *Gephyrosaurus bridensis*), and a stem-lepidosaurian (*Sophineta cracoviensis*). To provide a phylogenetic framework for our analyses, we assembled an informal super-tree^43^ for the 548 taxa. For topology we followed the total evidence phylogeny of Simões *et al.*^44^ – the first work to find agreement between morphological and molecular evidence regarding early squamate evolution. The same source provided time calibrations for *Sophineta cracoviensis*, fossil and extant Rhynchocephalia, stem squamates and crown squamate groups. Using additional sources, we gathered complementary information on the stem and crown of Gekkota^45-47^, Dibamia^47^, Scincoidea^46,47^, Lacertoidea^46-52^ including Polyglyphanodontia^46,53^, Mosasauria^46^, Serpentes^46,47^, Anguimorpha^46,47^, and Iguania^46,47,54-56^. To avoid over-sampling Liolaemidae, we randomly selected species according to relative abundance of dietary categories within the group^34^ and of liolaemids among squamates. In the absence of time-calibrated phylogenetic information, we used temporal ranges from the Paleobiology Database (http://www.paleobiodb.org) and checked accuracy by comparison with cited sources. Each squamate group stated above was grafted onto the backbone of the Simões *et al.*^44^ phylogeny according to its proposed calibrations. Node calibrations falling within the 95% highest posterior density for the corresponding node in the Simões *et al.*^44^ phylogeny were kept unchanged. Where a calibration fell beyond that range, the calibration of Simões *et al.*^44^ was preferred. For taxa and nodes not included in Simões *et al.*^44^ and with phylogenetic data lacking time-calibration, we used the code of Mitchell *et al.*^57,58^ to generate calibrations based on last appearance dates and estimated rates of speciation, extinction, and preservation. The method – derived from Bapst^59^ – allows the stochastic estimation of node age based on the inferred probability of sampling a fossil and probability density of unobserved evolutionary history, though nodes are sampled downwards towards the root rather than upwards from it. We used preliminary BAMM 2.6^57^ runs not including the taxa concerned to generate estimates of speciation and extinction rates and selected a preservation rate of 0.01 (see below). Our tree includes 27 unresolved nodes, denoting phylogenetic uncertainty. For methods requiring a fully dichotomous tree, we used the function multi2di in ape 5.3^60^ for R 3.6.1^61^ to generate a random dichotomous topology. Because of the sensitivity of BAMM to zero-length branches, we then used the method of Mitchell *et al*^58^. to generate non-zero branch lengths in randomly resolved polytomies with fossil taxa. We used the same randomly resolved and calibrated tree in all analyses requiring a dichotomous tree. We referred to the August 12^th^, 2019 version of the Reptile Database (http://www.reptile-database.org) and the Paleobiology Database (http://www.paleobiodb.org) for taxonomical reference of extant and fossil species (respectively).

### Dietary data

We followed Meiri^62^ and Pineda-Munoz & Alroy^63^ for dietary classification. Accordingly, when quantitative dietary data were available, we classified species based on the main feeding resource in adults (*i.e.*, >50% of total diet in volume^63^). Species consuming >50% plant material were classified as herbivores. We followed Meiri^62^ and Cooper & Vitt^64^ in defining omnivorous diets as including between 10 and 50% of plants, to account for accidental plant consumption by some predators. Among predators, carnivores are defined as feeding mostly on vertebrates. Predators consuming primarily arthropods and molluscs are “insectivores.” We could find no published dietary hypothesis for 64 out of 119 fossil species, which we assigned to the most plausible of our diet categories based on tooth complexity and the diets of closely related taxa (see Supplementary Data File 1).

### Geometric morphometrics

Specimens of 75 species were selected to represent all major groups of squamates with multiple-cusped teeth and based on the quality of the material available. We extracted two-dimensional outlines for geometric morphometric analyses from 52 X-ray computed microtomography scans (microCT-scans), 12 photographs, 10 anatomical drawings of specimens, and one scanning electron microscopy (SEM) image. Sources included the literature, the Digital Morphology (DigiMorph) library, four new photographs, and four new microCT-scans (see below and Supplementary Data File 1).

To analyse morphological variation of tooth shapes, we collected two-dimensional open outlines of a left upper posterior maxillary multicuspid tooth in labial view with ImageJ 1.47v^65^. We chose whenever possible the tooth with the most numerous cusps in the quadrant. If no left maxillary tooth was sampled or suitable for tracing an outline, we referred to the right quadrant or lower jaws and mirrored the outline adequately to retain the same orientation. We used the EqualSpace function from PollyMorphometrics 10.1^66^ for Mathematica 10^67^ to normalize teeth outlines as sets of 200 equally spaced points based on Bézier splines functions.

We used Momocs 1.3.0^68^ for R^61^ to perform geometric morphometric analyses of tooth outlines. We first applied a Bookstein alignment^69^ and, for each outline, computed by Discrete Cosine Transform (DCT) the first 21 harmonic amplitudes^70^. Harmonic coefficients were then processed by Principal Component Analysis (PCA)^71^. We limited graphical representation of the PCA to its first two axes, accounting for 89% of all morphological variation. To determine the significance of our dietary grouping, we fitted a phylogenetic multivariate linear model using penalized likelihood (PL)^72,73^ on all PC scores using mvMORPH 1.1.1^74^ for R^61^. Because we sampled only two carnivorous species, we added these to our insectivorous sample (n = 51, so making a “predatory” group, to avoid spurious conclusions arising from groups with extremely low sample sizes. Model fit was performed using Pagel’s λ^75^ to jointly estimate the phylogenetic signal in model residuals. We then used a one-way phylogenetic PL-MANOVA to evaluate overall differences between dietary groups. To test between-group differences, we used general linear hypothesis testing through contrast coding. We fitted a model for which each group was explicitly estimated to test compound contrasts.

### Ancestral character state reconstructions

We reconstructed the evolution of cusp number and diet using Maximum-Likelihood (ML) ancestral character state reconstruction under a time-reversible continuous Markov transition model^26,76^ as implemented in phytools 0.6-99^77^. We retrieved marginal ancestral states at the nodes of the tree with the re-rooting algorithm from the same package^78^ and generated a model of character evolution by averaging three character transition matrices (all transitions allowed with either all rates different, symmetrical rates, or equal rates) according to their respective fit through Akaike-weights model averaging^79^ (see Supplementary Table I). Finally, we used stochastic character mapping (1,000 simulations) to compute the most likely character states at each node based on the model-averaged transition matrix^80^. In contrast with tooth complexity ancestral states reconstructions, extant data allow the formulation of informed hypotheses on possible dietary transitions in squamates. Insects are an important food resource for the juveniles of many squamate species, and several extant species of plant consumers show an ontogenetic dietary shift from insectivorous juveniles to omnivorous or herbivorous adults^36,64,81,82^. Moreover, extant data show that predatory squamates may rely on plant material depending on environmental conditions^34,64,83-85^. Therefore, it has been hypothesized that squamate plant consumption originated in predatory animals, which evolved increasingly more plant-based diet through time under selective pressure^34,64^. We thus chose to test a specific hypothesis of dietary transitions against naïve models and base our reconstructions on the best-performing model. We compared the respective fit of three default models (all transitions allowed) to three variants of our hypothesis of dietary transitions (limiting transitions to carnivore-insectivore, insectivore-omnivore, and omnivore-herbivore, with three transition rate regimes) and selected the model with highest relative fit (*i.e.*, the custom model with all rates different) to retrieve ancestral states at the nodes (see Supplementary Table 1). Based on these ancestral reconstructions, we gathered a list of changes in cusp number and plant consumption. Subsequently, we identified pairs of increases or decreases in both traits belonging to the same phylogenetic path (the unique succession of branches connecting a descendent lineage to one of its ancestors) and noted whether each initiated by a change in cusp number, plant consumption, or whether both changes happened on the same branch.

### Tests of correlated evolution

We used BayesTraits 3.0.2 (www.evolution.rdg.ac.uk) to run Markov Chain Monte Carlo (MCMC) models of evolution of tooth complexity and diet with independent or correlated (*i.e.* assuming rates of transition in one trait depend on the character state of the other) rates of character transition. To improve rate estimations with our discrete dataset, in each run we scaled our tree to obtain an average branch length of 0.1 (*i.e.*, scaling factor = 5.017e-3) as recommended by the software manual. Due to method limitations, we transformed our discrete tooth complexity and diet characters into binary traits (teeth bearing one cusp vs two cusps or more, and carnivores and insectivores (predators) vs omnivores and herbivores (plant consumers), respectively). Each model ran for 110,000,000 iterations with default rate priors, and we discarded the first 10,000,000 iterations as burn-in. We sampled parameters every 10,000 iterations and checked each chain for convergence and large effective sample size (using CODA 0.19-3^86^ for R^61^). We used a steppingstone sampler^87^ to retrieve the marginal likelihood of each model (250 stones, each run for 10,000 iterations), which we compared with a log Bayes Factor to provide a measure of relative support of each model^88^. Analyses of the following combinations of different binarization of the dataset yielded similar results, *i.e.*, correlated evolution of cusp number and diet: one or two cusps vs three cusps or more combined with carnivores and insectivores vs omnivores and herbivores, and one to three cusps vs more than three cusps combined with herbivores vs other diets. As expected, a correlated model was weakly or not supported for other combinations (Supplementary Table 4).

### Rates of phenotypic evolution

We estimated the evolutionary rate of tooth shape change through the variable rates model of BayesTraits 3.0.2^89,90^. In this approach, a reversible-jump Markov Chain Monte Carlo (rjMCMC) algorithm is used to detect shifts in rates of continuous trait evolution – modelled by a Brownian motion (BM) process – across the branches of a phylogenetic tree. This is achieved by estimating the location of the shifts in rates (the product of a homogeneous background rate with a set of rate scalars) by using two different proposal mechanisms (one updating one branch at a time and one updating complete subclades). We used the default gamma priors on rate scalar parameters. Support for rate heterogeneity was then further confirmed by comparing the fit of the variable rates model against a null single-rate Brownian model. Here, we ran a variable rates model and a homogeneous Brownian model on the scores of the first 12 pPC axes from our phylogenetic PCA of tooth outlines, accounting for over 99% of the total variance. Because PC axes can be correlated in a phylogenetic context, we used the phylogenetic PC scores to remove evolutionary correlations^91,92^. We ran a phylogenetic principal component analysis (pPCA)^92^ on the first 21 harmonics obtained by DCT using phytools 0.6-99^77^ for R^61^. As for the original PCA, we found the two first pPC axes accounted for the largest part of all morphological variation (83% of cumulative variance). All parameters used were the same as for the correlated evolution tests (see above): 110,000,000 iterations, 10% burn in, default priors, rescaling factor = 5.017e-3, sampling every 10,000 iterations, convergence and sample size checks, stepping stone sampler with 250 stones run 10,000 iterations. Finally, we plotted our species tree (using phytools 0.6-99^77^, ggtree 1.8.1^93^, and viridis 0.5.1^94^ for R^61^) with branches scaled by the averaged rate scalars across posterior samples (returned by the Variable Rates Post Processor; http://www.evolution.reading.ac.uk/VarRatesWebPP/), thus indicating the relative deviation from the background rate of change.

Likewise, we used a variable rates model approach on discrete data to detect heterogeneity in character transition rates for tooth complexity and diet. The variable rates model operates on discrete data by breaking the assumption of a single character transition rate matrix defined for the entire tree, which it achieves by re-scaling this transition matrix in different parts of the tree using an rjMCMC algorithm. As for continuous data, the process generates a posterior distribution of scalars for each branch, and comparison with a null MCMC model with a constant transition matrix allows evaluation of support for heterogeneity in the strength of character transition rates. We ran the variable rates and null models similarly to tooth shape data (see above), using binarized tooth complexity and dietary data to avoid over-parameterization. The Variable Rates Post Processor returned the averaged branch rate scalars used to plot the tree according to local deviations from the background transition matrix. The large variances returned by the post-processor for some rate scalars, however, denote a relatively complex model to fit and warrant adequate caution in interpreting absolute rate scalar values, though relative rate differences should be fully representational. For clarity, we coloured each branch according to a common log-transformed scale. We again tested alternative binarizations of diet and tooth complexity and found support for a variable rates model for the alternative binarization of diet and one other binarization of tooth complexity (one or two cusps vs three cusps or more) (Supplementary Table 5).

### Models of diversification

We fitted different trait-dependent models of speciation and extinction (BiSSE, HiSSE) and associated trait-independent null models with hisse 1.9.5^95^ for R^61^, for which we compared relative fit using Akaike weights^79^. In addition, we used an rjMCMC algorithm with BAMM 2.6^57^ and BAMMtools 2.1.6 for R^61^ to model rates of speciation and extinction independently from trait evolution on a random resolution of our super-tree (see above). This is currently the only available method allowing branch-specific estimation of diversification rates on non-ultrametric trees (*i.e.*, including fossil taxa) by using a fossilized birth-death process^57^. We ran ten independent replicates for 20,000,000 generations using priors generated by the setBAMMpriors function of BAMMtools, a preservation rate prior of 0.01 (to reflect the sampling biases affecting the squamate fossil record^96^), and a global sampling fraction of 0.048 accounting for our sampling relative to the total diversity of living and extinct squamates referenced in both the Reptile Database (http://www.reptile-database.org) and the Paleobiology Database (http://www.paleobiodb.org). We set a 10% burn-in and checked convergence and effective sample size with CODA 0.19-3^86^. Because we encountered many equiprobable configurations, for each run we computed the maximum shift credibility (MSC) configuration and extracted speciation and extinction rates for clades defined by each node immediately above a shift, plus mean rates outside these clades (background rate). We then calculated a mean shift magnitude for each clade using the ratio of its mean speciation rate over the mean background rate^97^. To control for the influence of aquatic taxa during the KTR, we repeated analyses on a tree devoid of Cretaceous aquatic taxa (ten mosasaurs and three snakes) and found no changes to our results.

### Statistics

We performed all univariate non-parametric tests using rcompanion 2.3.25 (https://www.rcompanion.org) and the base stat package in R 3.6.1^61^. All effect sizes^98^ and their 95% confidence intervals were computed by bootstrap over 10,000 iterations. Sample size for all tests is n = 548. A Kruskal–Wallis *H* test^99^ on tooth complexity levels among squamate dietary categories showed a statistically significant effect of diet on the level of tooth complexity (*χ*^2^ = 144.27, df = 3, *p*-value = 4.5e-31, ε^2^ = 0.26 [0.20, 0.34]). Post-hoc pairwise two-sided Wilcoxon–Mann–Whitney tests^100,101^ showed statistically significant differences between all dietary categories (see Extended Data Table 1 for full reporting).

We used mvMORPH 1.1.1^74^ for R^61^ to perform regularised phylogenetic one-way multivariate analyses of variance (MANOVA) and multivariate general linear hypothesis tests in a penalized likelihood framework^72,73^. For each test, we assessed significance over 10,000 permutations of the Pillai trace^102^ obtained through regularised estimates^72,73^. A regularised phylogenetic MANOVA on the principal component scores of 75 tooth outlines showed statistically significant differences in 2D tooth shape between diets (V = 1.04, *p*-value = 0.001). We then used general linear hypothesis tests to evaluate simple and compound contrasts between groups, of which all but one were statistically significantly different (see Extended Data Table 2 for full reporting).

Two-sided Wilcoxon–Mann–Whitney tests^100,101^ on macroevolutionary rates inferred using the best-performing trait-dependent model of speciation and extinction (see Extended Data Table 4 and Figure 5) show multiple-cusped taxa have both statistically significantly higher speciation and extinction rates than taxa with single-cusped teeth. Likewise, plant-consuming (*i.e.*, omnivorous and herbivorous) taxa have both statistically significantly higher speciation and extinction rates than mainly predatory taxa (*i.e.*, carnivores and insectivores) (see Extended Data Figure 5 for full reporting).

### Photographs and X-ray computed microtomography

Photographs of ten specimens were captured at the Museum für Naturkunde (Berlin, Germany). New microCT-scan data was generated for 24 specimens using a Skyscan 1272 microCT (Bruker) at the University of Helsinki (Finland), a Skyscan 1172 microCT (Bruker) at the University of Eastern Finland (Kuopio, Finland), and a Phoenix nanotom CT (GE) at the Museum für Naturkunde (Berlin, Germany). Three-dimensional surface renderings were generated using Amira 5.5.0^103^.

### Specimen collection

Specialised retailers provided specimens of five species (see Supplementary Data File 1). The Laboratory Animal Center (LAC) of the University of Helsinki and/or the National Animal Experiment Board (ELLA) in Finland approved all reptile captive breeding (license numbers ESLH-2007-07445/ym-23 and ESAVI/7484/04.10.07/2016).

### Art credits

Figure 1: the authors after Darren Naish (used with permission), Phylopic courtesy of Michael Keesey, David Orr, Ian Reid, Alex Slavenko, and Steven Traver, and public domain. Figure 2: the authors after Dick Culbert (CC-BY 2.0), Scott Robert Ladd (CC-BY 3.0), and Darren Naish (used with permission), Phylopic courtesy of Michael Croggie, Michael Keesey, Alex Slavenko, and Jack Meyer Wood. See https://www.phylopic.org for additional license information.

### Data availability statement

All datasets generated and analysed during the current study (tip-state dataset, polytomous and dichotomous versions of our phylogeny, 2D outlines; see Fig. 1-4 and Extended Data Figure 1-5) are available as Supplementary Information. CT-scan data are available through NDP, upon reasonable request.

## Acknowledgements

We thank Ilpo Hanski and Martti Hildén (Luonnontieteellinen keskusmuseo, Helsinki, Finland) for specimen loans, Johannes Müller (Museum für Naturkunde, Berlin, Germany) for specimen loans and access to collections and CT-scanning facilities, Jessie Maisano (University of Texas, Austin, TX) for sharing data from the DigiMorph database, Arto Koistinen (University of Kuopio, Finland) and Heikki Suhonen (University of Helsinki, Finland) for access to CT-scanning facilities, Arto Koistinen, Simone Macrì, Kristin Mahlow, and Filipe Oliveira da Silva for acquiring morphological data, as well as Jukka Jernvall, Mikael Fortelius, and the Helsinki Evo-Devo community for discussions. We thank Vincent Bonhomme, David Caetano, Andrew Meade, and Johnathan Mitchell for their help in implementing Momocs, HiSSE models, BayesTraits, and BAMM 2.6 respectively. We also thank Robert Espinoza for precisions on liolaemid diets. This work was supported by funds from the Integrative Life Science doctoral program (ILS; to FL), the Center for International Mobility scholarship program (CIMO; to FL), the University of Helsinki (to NDP), the Institute of Biotechnology (to NDP), Biocentrum Helsinki (to NDP), and the Academy of Finland (to NDP).

## Authors contributions

FL, IJC, and NDP designed the experimental approach. FL and NDP collected the specimens for microCT-scanning. FL character-coded species from the literature and specimen data. FL collected tooth outline semi-landmark data. FL performed the research. FL analysed the data, with contribution from JC, IJC and NDP. FL made the figures. FL produced the first draft, and FL, JC, and IJC wrote the paper, to which all authors contributed in the form of discussion and critical comments. All authors approved the final version of the manuscript.

## Competing interest declaration

The authors declare no conflict of interests.

## Additional information

Correspondence and requests for materials should be addressed to FL, IJC, and NDP. Supplementary information is available for this paper.

---

**Extended Data Figure 1.**
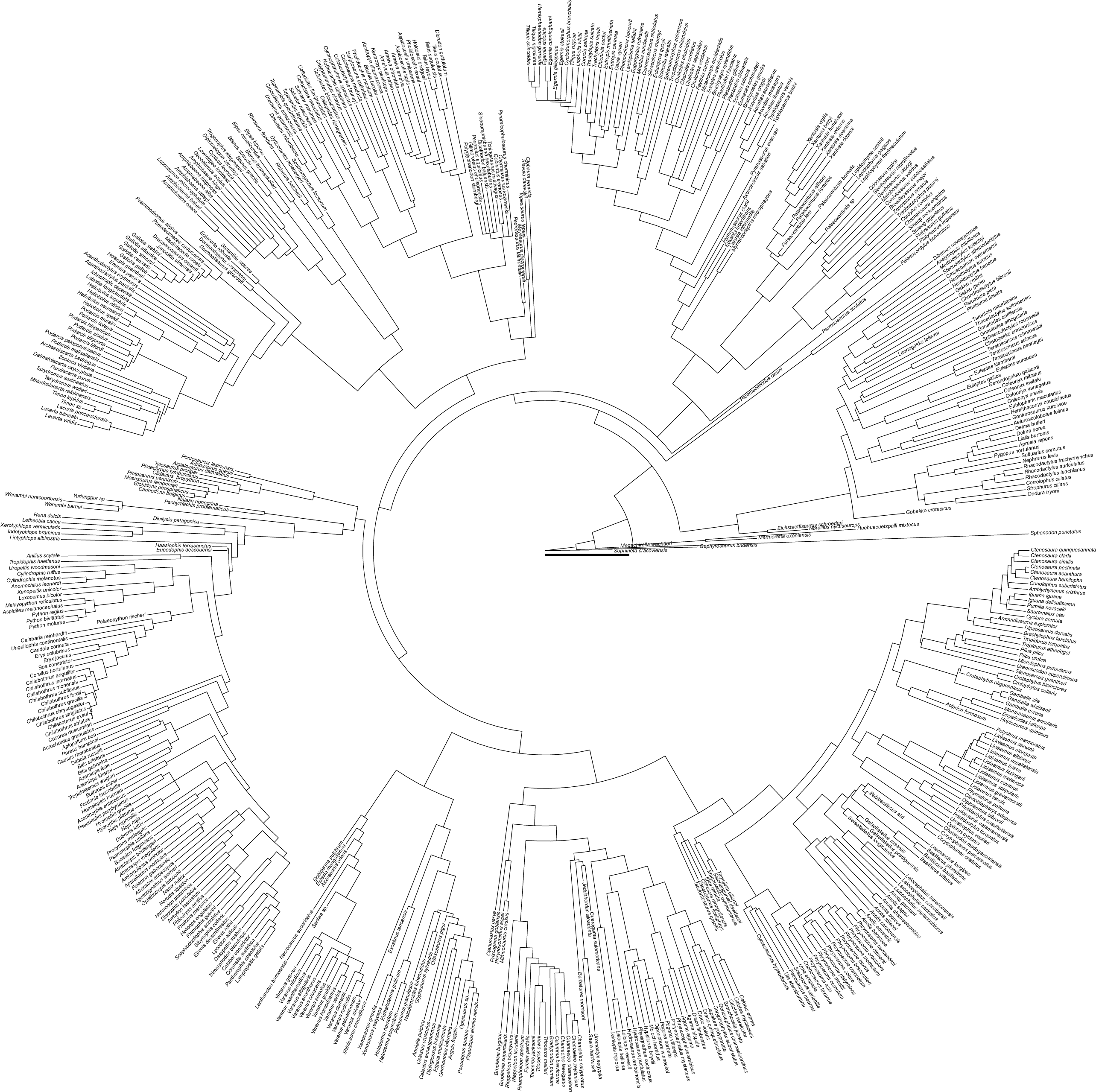
Time-calibrated squamate phylogeny. Informal super-tree including 545 extant and extinct squamates species and three outgroup species (see Methods). Scalebar = 50 million years.

**Extended Data Figure 2.**
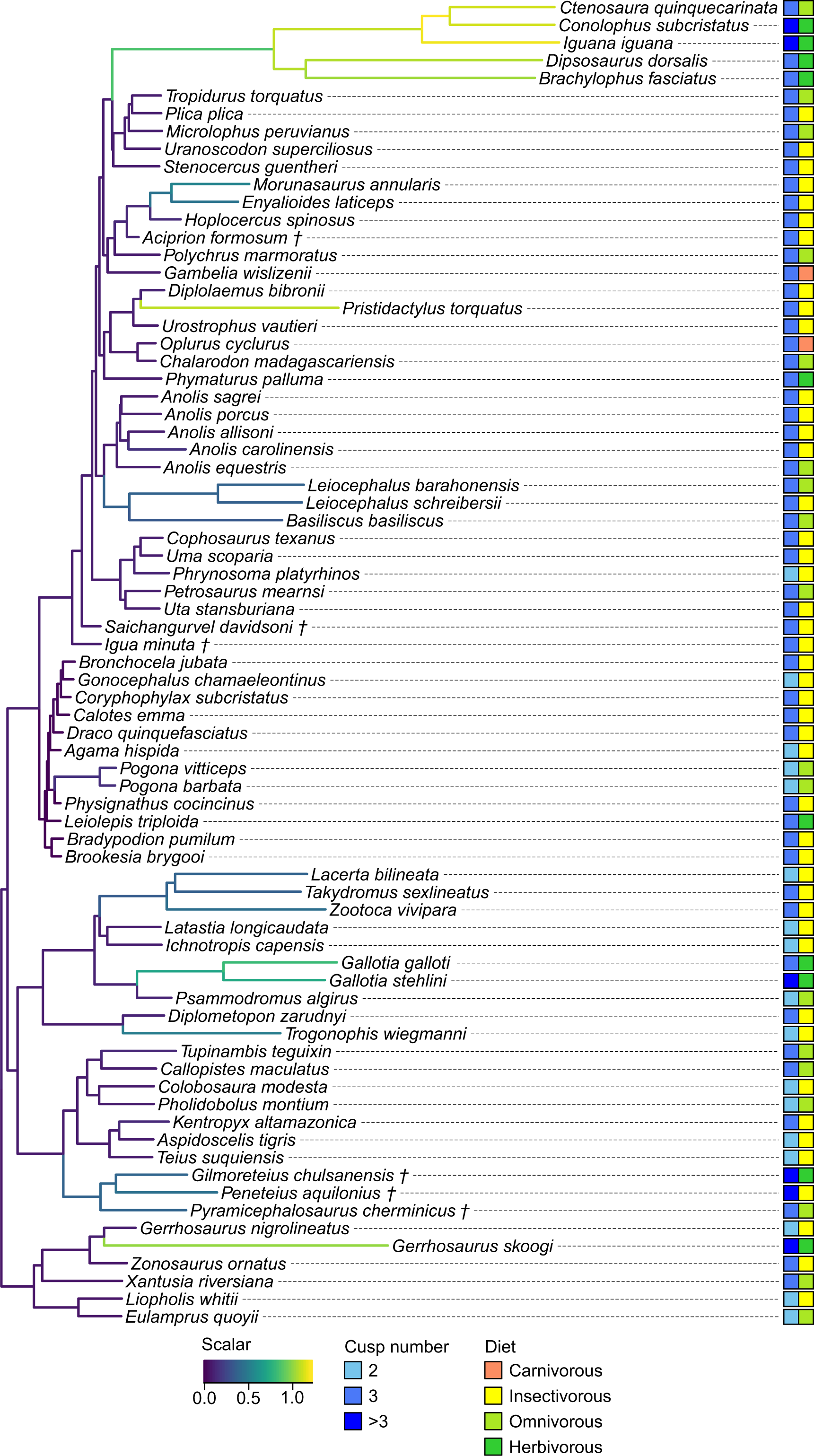
Rates of 2D tooth shape evolution among 75 squamate species with multicuspid teeth. Branch lengths are transformed by the mean of the respective posterior distribution of scalars generated under a variable rates model, reflecting changes in the rate of shape evolution. † = extinct taxon.

**Extended Data Figure 3.**
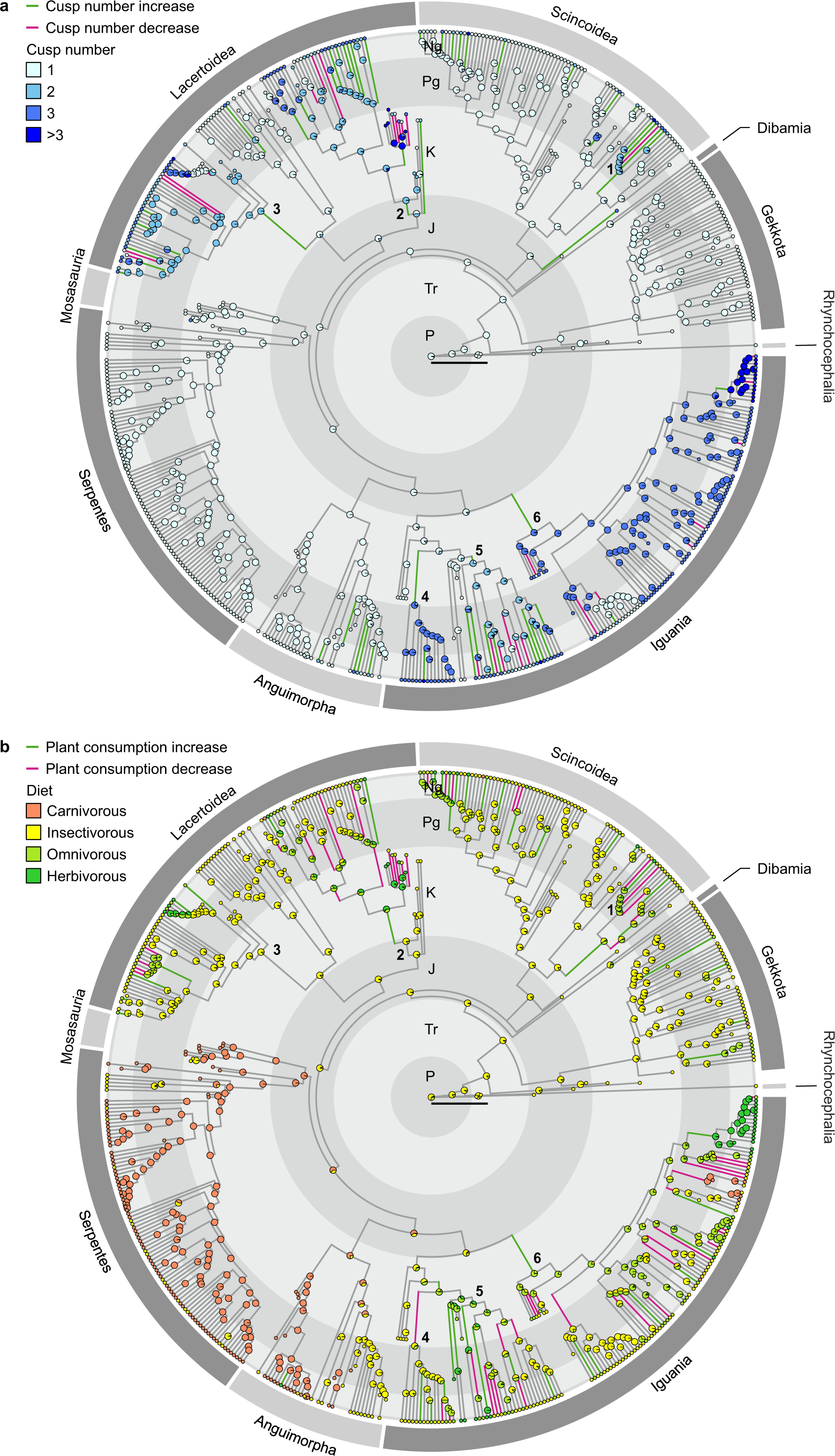
Squamate dental and dietary evolution. Known and Maximum-Likelihood ancestral state reconstructions of tooth complexity (**a**) and diet (**b**) in squamates. Pie charts indicate the relative likelihood of each character state at the corresponding node. Branch tip circles indicate character state at tips. Coloured branches indicate an increase or a decrease in tooth complexity/plant consumption. 1: Gerrhosauridae. 2: Teiioidea + Polyglyphanodontia (informally Teiioidea *sensu lato*). 3: total group Lacertidae (informally Lacertidae *sensu lato*). 4: Chamaeleonidae. 5: non-Uromastycinae agamids (informally Agamidae *sensu stricto*). 6: total group Pleurodonta. P: Permian. Tr: Triassic. J: Jurassic. K: Cretaceous. Pg: Paleogene. Ng: Neogene. Scalebar = 50 million years.

**Extended Data Figure 4.**
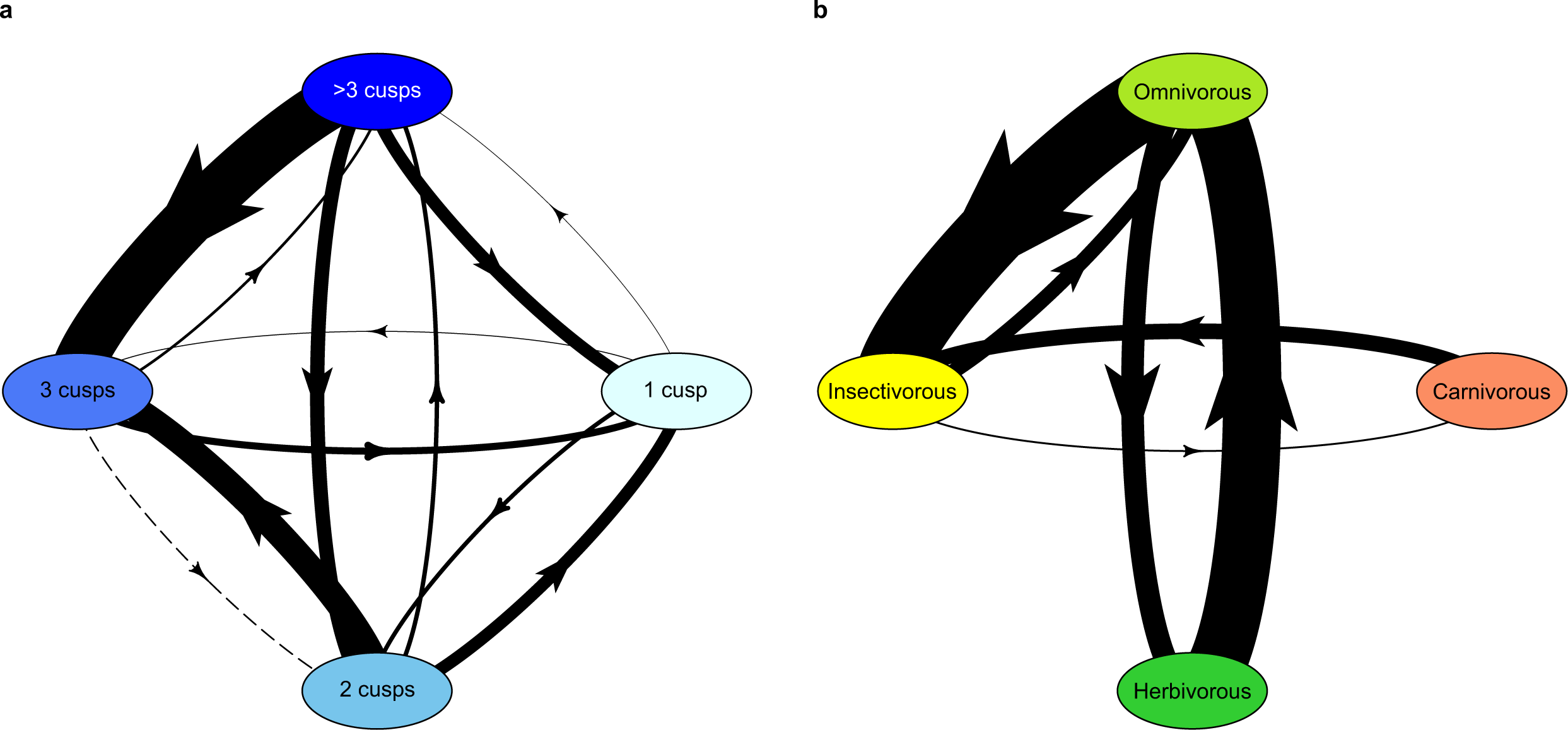
Transition models used for ancestral state reconstructions. Relative character transition rates for tooth complexity (**a**) and diet (**b**). Arrow widths are scaled by the log-transformed rates of transition. The orientation of arrows denotes the direction of character transitions. Note: the transition from two-cusped to three-cusped teeth is represented with a dotted line, due to its relative rate being negligible compared to all other transition rates.

**Extended Data Figure 5.**
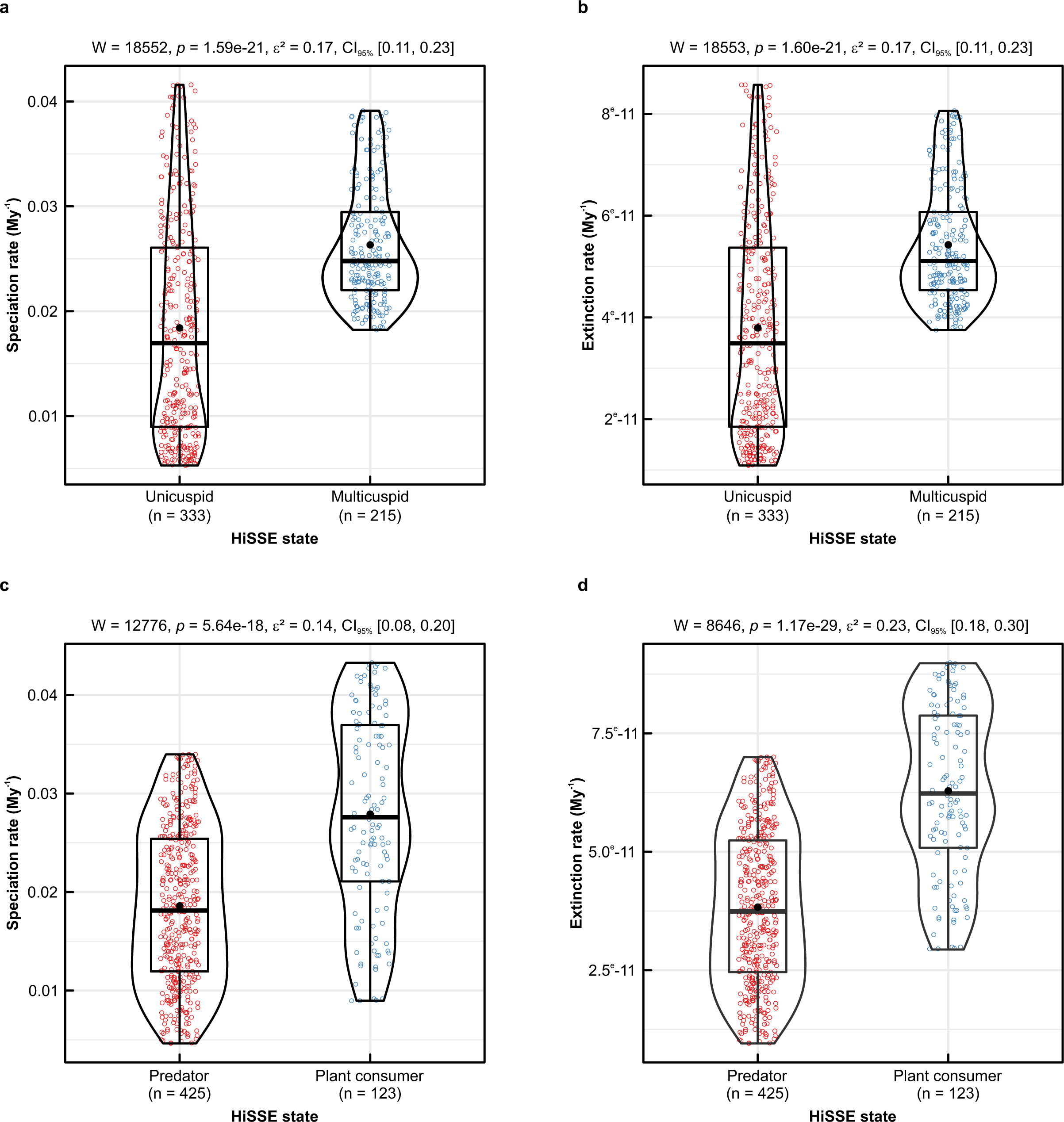
Squamate macroevolutionary rates among different levels of tooth complexity and diet. Speciation (**a–c**) and extinction rates (**b–d**) per tooth complexity or diet character state for the best supported model of trait-dependent speciation and extinction. Violin plots indicate the density of data points. Boxes include 50% of the data points, with the black line and dot indicating the median and mean, respectively. Whiskers incorporate the whole range of the data. All pairs are statistically significantly different (Wilcoxon–Mann– Whitney test); see panels for their respective W statistic and effect size (ε^2^), including 2.5^th^ and 97.5^th^ confidence interval percentiles (CI_95%_) in brackets.

**Extended Data Table 1.** Tooth complexity of squamates analysed by dietary category.

| Group 1 | Group 2 | Sample size | W | p-value |
| --- | --- | --- | --- | --- |
| Carnivores | Herbivores | 154 | 490 | **4.9e-21** |
| Carnivores | Insectivores | 425 | 12458 | **1.8e-08** |
| Carnivores | Omnivores | 197 | 1583 | **7.3e-21** |
| Herbivores | Insectivores | 351 | 10188 | **8.2e-13** |
| Herbivores | Omnivores | 123 | 2330 | **4.6e-04** |
| Insectivores | Omnivores | 394 | 7543 | **4.5e-10** |

Two-sided pairwise Wilcoxon–Mann–Whitney tests on the tooth complexity level of 548 species grouped by diet (carnivores, $n = 114$ ; insectivores, $n = 311$ ; omnivores, $n = 83$ ; herbivores, $n = 40$ ), with $W$ statistic and Bonferroni-corrected $p$ values. Significant differences ( $p < 0.05$ ) are in bold.

**Extended Data Table 2.** Pairwise comparisons of dietary categories in 2D tooth morphospace.

| Group 1 | Group 2 | Sample size | V | p-value |
| --- | --- | --- | --- | --- |
| Herbivores | Omnivores | 22 | 0.63 | **3.0<sup>e-04</sup>** |
| Herbivores | Predators | 64 | 0.76 | **1.0<sup>e-04</sup>** |
| Omnivores | Predators | 64 | 0.29 | 0.80 |
| Herbivores +<br>Omnivores | Predators | 75 | 0.64 | **2.0<sup>e-04</sup>** |
| Herbivores | Omnivores +<br>Predators | 75 | 0.73 | **1.0<sup>e-04</sup>** |

Multivariate general linear hypothesis testing through phylogenetic penalized likelihood MANOVA of PC scores for the multicuspid teeth of 75 extant and fossil squamate species grouped by diet (predators, n = 53 (including insectivores, n = 51, and carnivores, n = 2); omnivores, n = 11; herbivores, n = 11). We assessed statistical significance over 10,000 permutations of the Pillai trace (V). Significant differences ( $p < 0.05$ ) are in bold.

**Extended Data Table 3.** Squamate clades showing rate shifts in trait-independent models of speciation and extinction.

| Node | Replicates | Mean speciation rate (My<sup>-1</sup>) | Standard deviation | Ratio to mean outgroup rate | Mean extinction rate (My<sup>-1</sup>) | Standard deviation | Ratio to mean outgroup rate | Cusp number transition |
| --- | --- | --- | --- | --- | --- | --- | --- | --- |
| 555 (A) | 9/10 | 0.0523 | 0.0159 | 4.40 | 0.0092 | 0.0030 | 0.65 | - |
| 679 (B) | 6/10 | 0.1296 | 0.0659 | 2.59 | 0.0128 | 0.0083 | 1.36 | - |
| 691 | 1/10 | 0.0511 | 0.0207 | 1.01 | 0.0122 | 0.0049 | 1.82 | - |
| 697 (C) | 9/10 | 0.0875 | 0.0489 | 1.72 | 0.0961 | 0.0572 | 11.06 | 2 → >3 |
| 705 | 1/10 | 0.0523 | 0.0212 | 1.03 | 0.0004 | 0.0002 | 0.05 | - |
| 763 | 2/10 | 0.1918 | 0.0899 | 3.69 | 0.1133 | 0.0553 | 11.00 | 2 → 1 |
| 782 (D) | 10/10 | 0.1439 | 0.0482 | 2.86 | 0.0308 | 0.0247 | 3.37 | - |
| 802 (E) | 10/10 | 0.0596 | 0.0203 | 1.17 | 0.0557 | 0.0189 | 6.39 | - |
| 824 (F) | 7/10 | 0.0624 | 0.0296 | 1.26 | 0.0021 | 0.0014 | 0.21 | - |
| 826 | 1/10 | 0.0778 | 0.0315 | 1.54 | 0.0060 | 0.0024 | 0.49 | - |
| 846 (G) | 10/10 | 0.3912 | 0.0966 | 7.79 | 0.0433 | 0.0456 | 4.69 | - |
| 914 (H) | 10/10 | 0.1460 | 0.0231 | 2.92 | 0.0143 | 0.0111 | 1.55 | - |
| 996 (I) | 5/10 | 0.0797 | 0.0417 | 1.71 | 0.0100 | 0.0052 | 1.15 | 1 → 3 |
| 1003 (J) | 5/10 | 0.0802 | 0.0420 | 1.71 | 0.0056 | 0.0030 | 0.61 | - |
| 1010 (K) | 10/10 | 0.1586 | 0.0154 | 3.16 | 0.0159 | 0.0153 | 1.72 | 3 → 1 |
| 1055 (L) | 10/10 | 0.3224 | 0.0939 | 6.43 | 0.0268 | 0.0181 | 2.89 | - |
| 1084 (M) | 6/10 | 0.2555 | 0.1330 | 5.15 | 0.0272 | 0.0198 | 3.04 | 3 → >3 |
| 1085 | 4/10 | 0.2934 | 0.1555 | 5.91 | 0.0347 | 0.0260 | 3.63 | - |

**Extended Data Table 4.** Tests of trait-dependent models of squamate speciation and extinction.

| Model | Hidden states | Character-dependent | Number of diversification rates | Number of transition rates | Dual transitions | AICc (tooth complexity) | ωAICc (tooth complexity) | AICc (diet) | ωAICc (diet) |
| --- | --- | --- | --- | --- | --- | --- | --- | --- | --- |
| HiSSE complete ARD | Yes | Yes | 4 | 12 | Yes | 5231.30 | 0.004 | 5301.83 | 0.068 |
| HiSSE complete ER | Yes | Yes | 4 | 1 | Yes | 5275.11 | < 0.001 | 5381.02 | < 0.001 |
| **HiSSE node dual ARD** | **Yes** | **Yes** | **4** | **8** | **No** | **5221.05** | **0.595** | **5296.89** | **0.802** |
| HiSSE no dual 3 rates | Yes | Yes | 4 | 3 | No | 5247.47 | < 0.001 | 5306.42 | 0.007 |
| HiSSE no dual ER | Yes | Yes | 4 | 1 | No | 5271.91 | < 0.001 | 5370.23 | < 0.001 |
| BiSSE | No | Yes | 2 | 2 | - | 5285.85 | < 0.001 | 5362.89 | < 0.001 |
| CID-4 3 rates | Yes | No | 4 | 3 | No | 5221.96 | 0.378 | 5300.65 | 0.123 |
| CID-4 ER | Yes | No | 4 | 1 | No | 5244.80 | < 0.001 | 5353.53 | < 0.001 |
| CID-2 complete ARD | Yes | No | 2 | 12 | Yes | 5245.97 | < 0.001 | 5362.36 | < 0.001 |
| CID-2 complete ER | Yes | No | 2 | 1 | Yes | 5270.88 | < 0.001 | 5377.14 | < 0.001 |
| CID-2 no dual ARD | Yes | No | 2 | 8 | No | 5239.53 | < 0.001 | 5345.35 | < 0.001 |
| CID-2 no dual 3 rates | Yes | No | 2 | 3 | No | 5227.58 | 0.023 | 5310.76 | < 0.001 |
| CID-2 no dual ER | Yes | No | 2 | 1 | No | 5260.71 | < 0.001 | 5364.75 | < 0.001 |

Description of model parameters and relative goodness of fit. For both tooth complexity and plant consumption, a “hidden state” speciation and extinction model (HiSSE) is best supported to account for squamate macroevolutionary patterns. Corrected Akaike Information Criterion (AICc) values and AICc model weights ( $\omega$ AICc) for the binary tooth complexity dataset and the binary diet dataset. The model with the highest AICc weight is highlighted in bold. HiSSE: hidden state speciation and extinction model; BiSSE: binary state speciation and extinction model; CID: character-independent (null) model; “complete”: all character state transitions possible, including simultaneous transitions in both the observed and hidden trait; “no dual”: simultaneous transitions in both the observed and hidden trait are excluded; ARD: all rates different; ER: equal rates.

## Notes

### Competing Interest Statement

The authors have declared no competing interest.

