## Supplementary Tables for "Multiple evolutionary origins and losses of tooth complexity in squamates"

This study is supported by the following supplementary information:

**Supplementary Tables:** includes Supplementary Tables 1–6, covering additional details of ancestral character state reconstructions, tests of correlated evolution, rates of phenotypic evolution, and rates of speciation and extinction models.

**Supplementary Data 1:** a time-calibrated phylogenetic tree of 545 squamate species and their outgroup, in Newick format.

**Supplementary Data 2:** a dichotomous time-calibrated phylogenetic tree of 545 squamate species and their outgroup, in Newick format.

**Supplementary Data 3:** 75 two-dimensional tooth outlines in an archive file. File names are formatted for use with Momocs 1.1.1 for R (*i.e.*, "Genus-species_Diet_status.txt").

**Supplementary Data 4:** species-level dataset in CSV format including taxonomic information (columns "species", "order", "suborder", "family"), living/fossil status ("status"), four-state tooth complexity level ("tooth.complexity") with three alternative binarization ("tooth.complexity.bin1", "tooth.complexity.bin2", "tooth.complexity.bin3"), diet data ("diet") with two alternative binarizations ("diet.bin1" and "diet.bin2"), references for tooth complexity, including a unique identifier and specimen identification ("reference.teeth", "reference.teeth.id", "specimen"), references for diet data with a unique identifier ("reference.diet", "reference.diet.id"), rate scalars for the variable rates model of tooth complexity and diet ("var.rates.teeth", "var.rates.diet"), speciation and extinction rates averaged over ten independent BAMM replicates ("mean.spe", "mean.ext"), speciation and extinction rates for the best performing tooth complexity- and diet-dependent HiSSE model ("hisse.teeth.spe", "hisse.teeth.ext", "hisse.diet.spe", "hisse.diet.ext").

### Supplementary Tables

#### Supplementary Table 1 | Model selection for ancestral character states reconstructions.

| Model | Number of transitions | Number of rates | AICc (tooth complexity) | $\omega$AICc (tooth complexity) | AICc (diet) | $\omega$AICc (diet) |
|---|---|---|---|---|---|---|
| **ARD** | **12** | **12** | **661.584** | **~1** | 815.412 | ~0.375 |
| SYM | 12 | 6 | 710.564 | < 0.001 | 877.410 | < 0.001 |
| ER | 12 | 1 | 737.734 | < 0.001 | 928.694 | < 0.001 |
| **progARD** | **6** | **6** | NA | NA | **814.387** | **~0.625** |
| progSYM | 6 | 3 | NA | NA | 877.601 | < 0.001 |
| progER | 6 | 1 | NA | NA | 1018.540 | < 0.001 |

Description of model parameters and relative goodness of fit. Corrected Akaike Information Criterion (AICc) values and AICc model weights ($\omega$AICc) for the binary tooth complexity dataset and the binary diet dataset. The model with the highest AICc weight is highlighted in bold. ARD: all rates different. SYM: symmetric rates (forces rate[A→B] = rate[B→A]). ER: equal rates. "prog" models correspond to the three rate variants of the dietary evolution model tested in this study.

**Supplementary Table 2 | Transition models used for the stochastic character mapping of cusp number and diet.**

| Character | q12 | q13 | q14 | q21 | q23 | q24 | q31 | q32 | q34 | q41 | q42 | q43 |
|---|---|---|---|---|---|---|---|---|---|---|---|---|
| Cusp number | 1.7$^{e}$-03 | 4.5$^{e}$-04 | 1.3$^{e}$-04 | 5.2$^{e}$-03 | 1.1$^{e}$-02 | 1.9$^{e}$-03 | 3.0$^{e}$-03 | 7.6$^{e}$-14 | 1.1$^{e}$-03 | 5.4$^{e}$-03 | 6.1$^{e}$-03 | 2.2$^{e}$-02 |
| Diet | 6.6$^{e}$-03 | 0 | 0 | 8.5$^{e}$-04 | 6.3$^{e}$-03 | 0 | 0 | 3.0$^{e}$-02 | 9.6$^{e}$-03 | 0 | 0 | 2.5$^{e}$-02 |

q: relative transition rate between character states. Character states are encoded as follows: Cusp number (1 – single-cusped, 2 – two-cusped, 3 – three-cusped, 4 – more than three cusps); Diet (1 – carnivore, 2 – insectivore, 3 – omnivore, 4 – herbivore).

#### Supplementary Table 3 | Events of paired increases or decreases in tooth complexity and plant consumption.

| Tooth complexity transition | Branch | Diet transition | Branch | Transition type | Pair type |
|---|---|---|---|---|---|
| 1 cusp to 2 cusps | 132 | insectivore to omnivore | 118 | increase | plant consumption first |
| 2 cusps to 1 cusp | 139 | omnivore to insectivore | 139 | decrease | co-occurrent |
| 2 cusps to >3 cusps | 143 | omnivore to herbivore | 143 | increase | co-occurrent |
| 1 cusp to 2 cusps | 154 | insectivore to herbivore | 154 | increase | co-occurrent |
| 1 cusp to 3 cusps | 167 | insectivore to omnivore | 167 | increase | co-occurrent |
| 1 cusp to 2 cusps | 230 | insectivore to omnivore | 230 | increase | co-occurrent |
| 1 cusp to >3 cusps | 250 | insectivore to herbivore | 250 | increase | co-occurrent |
| 1 cusp to 3 cusps | 263 | omnivore to herbivore | 263 | increase | co-occurrent |
| 1 cusp to 2 cusps | 272 | insectivore to omnivore | 254 | increase | plant consumption first |
| 1 cusp to 2 cusps | 280 | insectivore to herbivore | 284 | increase | tooth complexity first |
| 2 cusps to >3 cusps | 285 | insectivore to herbivore | 284 | increase | plant consumption first |
| >3 cusps to 3 cusps | 286 | herbivore to omnivore | 286 | decrease | co-occurrent |
| >3 cusps to 1 cusp | 289 | herbivore to insectivore | 289 | decrease | co-occurrent |
| >3 cusps to 1 cusp | 292 | herbivore to insectivore | 292 | decrease | co-occurrent |
| >3 cusps to 2 cusps | 293 | herbivore to omnivore | 293 | decrease | co-occurrent |
| 2 cusps to 3 cusps | 299 | omnivore to herbivore | 299 | increase | co-occurrent |
| 2 cusps to 1 cusp | 326 | omnivore to insectivore | 326 | decrease | co-occurrent |
| 2 cusps to 1 cusp | 330 | omnivore to insectivore | 328 | decrease | plant consumption first |
| 2 cusps to 1 cusp | 334 | omnivore to insectivore | 328 | decrease | plant consumption first |
| 2 cusps to 3 cusps | 347 | omnivore to herbivore | 354 | increase | tooth complexity first |
| 3 cusps to 1 cusp | 357 | omnivore to insectivore | 355 | decrease | plant consumption first |
| 1 cusp to 2 cusps | 398 | insectivore to omnivore | 408 | increase | tooth complexity first |
| 1 cusp to 2 cusps | 398 | insectivore to omnivore | 447 | increase | tooth complexity first |
| 1 cusp to >3 cusps | 417 | insectivore to omnivore | 417 | increase | co-occurrent |
| 1 cusp to >3 cusps | 417 | omnivore to herbivore | 418 | increase | tooth complexity first |
| >3 cusps to 3 cusps | 419 | herbivore to omnivore | 424 | decrease | tooth complexity first |
| 2 cusps to 3 cusps | 463 | insectivore to omnivore | 464 | increase | tooth complexity first |
| 2 cusps to 3 cusps | 475 | insectivore to omnivore | 478 | increase | tooth complexity first |
| 2 cusps to 3 cusps | 484 | insectivore to omnivore | 484 | increase | co-occurrent |
| 1 cusp to 3 cusps | 783 | insectivore to omnivore | 794 | increase | tooth complexity first |
| 1 cusp to 3 cusps | 783 | insectivore to omnivore | 800 | increase | tooth complexity first |
| 1 cusp to 3 cusps | 783 | insectivore to omnivore | 805 | increase | tooth complexity first |
| 1 cusp to 2 cusps | 822 | omnivore to herbivore | 823 | increase | tooth complexity first |
| 1 cusp to 2 cusps | 822 | insectivore to omnivore | 782 | increase | plant consumption first |
| 2 cusps to 3 cusps | 824 | omnivore to herbivore | 823 | increase | plant consumption first |
| 2 cusps to 3 cusps | 825 | omnivore to herbivore | 823 | increase | plant consumption first |
| 3 cusps to 1 cusp | 826 | herbivore to omnivore | 825 | decrease | plant consumption first |
| 2 cusps to 3 cusps | 832 | omnivore to herbivore | 830 | increase | plant consumption first |

| | | | | | |
|---|---|---|---|---|---|
| 2 cusps to 1 cusp | 837 | omnivore to insectivore | 837 | decrease | co-occurrent |
| 2 cusps to 1 cusp | 846 | omnivore to insectivore | 845 | decrease | plant consumption first |
| 2 cusps to 1 cusp | 854 | omnivore to insectivore | 852 | decrease | plant consumption first |
| 2 cusps to 3 cusps | 864 | insectivore to omnivore | 865 | increase | tooth complexity first |
| 1 cusp to 3 cusps | 869 | insectivore to omnivore | 869 | increase | co-occurrent |
| 1 cusp to 3 cusps | 869 | omnivore to herbivore | 1037 | increase | tooth complexity first |
| 3 cusps to 1 cusp | 874 | omnivore to insectivore | 874 | decrease | co-occurrent |
| 3 cusps to 1 cusp | 890 | omnivore to insectivore | 883 | decrease | plant consumption first |
| 3 cusps to 1 cusp | 895 | omnivore to insectivore | 883 | decrease | plant consumption first |
| 3 cusps to 1 cusp | 959 | omnivore to insectivore | 954 | decrease | plant consumption first |
| 3 cusps to 1 cusp | 972 | omnivore to insectivore | 967 | decrease | plant consumption first |
| 3 cusps to 1 cusp | 1021 | omnivore to carnivore | 1012 | decrease | plant consumption first |
| 3 cusps to >3 cusps | 1043 | omnivore to herbivore | 1037 | increase | plant consumption first |
| >3 cusps to 3 cusps | 1067 | herbivore to omnivore | 1067 | decrease | co-occurrent |

Transition locations are referred to by their branch number.

**Supplementary Table 4 | Tests of correlated and uncorrelated models of evolution of tooth complexity and diet in Squamata.**

| Tooth complexity binarization | Diet binarization | Independent model logLik | Dependent model logLik | logBF |
|---|---|---|---|---|
| One cusp vs two cusps or more | Omnivores and herbivores vs carnivores and insectivores | -446.16 | -435.74 | **20.85** |
| One cusp vs two cusps or more | Herbivores vs other diets | -317.08 | -317.97 | -1.78 |
| One or two cusps vs three or more | Omnivores and herbivores vs carnivores and insectivores | -432.12 | -417.81 | **28.62** |
| One or two cusps vs three or more | Herbivores vs other diets | -303.18 | -301.51 | 3.32 |
| Three cusps or less vs four or more | Omnivores and herbivores vs carnivores and insectivores | -303.15 | -304.83 | -3.37 |
| Three cusps or less vs four or more | Herbivores vs other diets | -174.33 | -162.91 | **22.84** |

Marginal log-likelihoods (logLik) for independent and dependent models of character evolution, with their associated log Bayes Factor (logBF), over three binarizations of tooth complexity level, two binarizations of diet, and their combinations. Values in bold (logBF > 10) indicate very strong statistical support for the more complex model (*i.e.*, dependent model).

**Supplementary Table 5 | Test of heterogeneity in relative character transition rates of tooth complexity and diet.**

| Character binarization | Constant model logLik | Variable rates model logLik | logBF |
|---|---|---|---|
| One cusp vs two cusps or more | -203.86 | -198.08 | **11.56** |
| One or two cusps vs three or more | -189.93 | -178.27 | **23.31** |
| Three cusps or less vs four or more | -61.29 | -62.36 | -2.14 |
| Omnivores and herbivores vs other diets | -241.64 | -232.49 | **18.30** |
| Herbivores vs other diets | -112.72 | -108.53 | 8.37 |

Marginal log-likelihoods (logLik) for constant and "variable rates" models of character evolution (allowing local heterogeneity in the character transition rates), with their associated log Bayes Factor (logBF), over three binarizations of tooth complexity level and two of diet. Values in bold (logBF > 10) indicate very strong statistical support for the more complex model (*i.e.*, variable rates model).

**Supplementary Table 6 | Tooth complexity and plant consumption changes in the vicinity of rate shifts in trait-independent models of speciation and extinction**

| Node | Cusp number transition (one node below) | Cusp number transition (one node above) | Plant consumption transition (one node below) | Plant consumption transition (one node above) |
|---|---|---|---|---|
| 555 (A) | - | - | - | - |
| 679 (B) | - | - | - | - |
| 691 | - | 1 → 2 | - | - |
| 697 (C) | - | >3 → 3 | I → H | H → O |
| 705 | - | - | I → H | - |
| 763 | - | - | - | - |
| 782 (D) | - | - | - | - |
| 802 (E) | - | - | - | - |
| 824 (F) | - | - | - | - |
| 826 | - | - | - | - |
| 846 (G) | - | - | - | - |
| 914 (H) | - | - | - | - |
| 996 (I) | - | - | - | - |
| 1003 (J) | - | - | - | O → I |
| 1010 (K) | - | - | - | - |
| 1055 (L) | - | - | - | O → I |
| 1084 (M) | - | - | O → H | - |
| 1085 | 3 → >3 | - | - | - |

All 18 clades defined by a rate shift in 10 maximum shift credibility configuration (MSC) independent replicates, including the 13 clades with rate shifts in at least five replicates (as indicated in Fig. 4c). Shift location is given by the number of the node immediately above it. The columns "Cusp number transition (one node below/above)" indicate changes in tooth complexity inferred in nodes immediately adjacent to a shift location, if any. Similarly, the columns "Plant consumption transition (one node below/above)" indicate changes in plant matter proportion in the diet in nodes immediately adjacent to a shift location, if any (I: insectivorous, O: omnivorous, H: herbivorous).
